# INSCT: Integrating millions of single cells using batch-aware triplet neural networks

**DOI:** 10.1101/2020.05.16.100024

**Authors:** Lukas M. Simon, Yin-Ying Wang, Zhongming Zhao

**Affiliations:** Center for Precision Health, School of Biomedical Informatics, The University of Texas Health Science Center at Houston, Houston, TX 77030, USA; Baylor College of Medicine, Therapeutic Innovation Center, Houston, TX, 77030, USA; Human Genetics Center, School of Public Health, The University of Texas Health Science Center at Houston, Houston, TX 77030, USA; MD Anderson Cancer Center UTHealth Graduate School of Biomedical Sciences, Houston, TX 77030, USA

## Abstract

Efficient integration of heterogeneous and increasingly large single cell RNA sequencing (scRNA-seq) data poses a major challenge for analysis and in particular, comprehensive atlasing efforts. Here, we developed a novel deep learning algorithm to overcome batch effects using batch-aware triplet neural networks, called INSCT (“Insight”). Using simulated and real data, we demonstrate that INSCT generates an embedding space which accurately integrates cells across experiments, platforms and species. Our benchmark comparisons with current state-of-the-art scRNA-seq integration methods revealed that INSCT outperforms competing methods in scalability while achieving comparable accuracies. Moreover, using INSCT in semi-supervised mode enables users to classify unlabeled cells by projecting them into a reference collection of annotated cells. To demonstrate scalability, we applied INSCT to integrate more than 2.6 million transcriptomes from four independent studies of mouse brains in less than 1.5 hours using less than 25 gigabytes of memory. This feature empowers researchers to perform atlasing scale data integration in a typical desktop computer environment. INSCT is freely available at https://github.com/lkmklsmn/insct.

**Highlights:**

- INSCT accurately integrates multiple scRNA-seq datasets
- INSCT accurately predicts cell types for an independent scRNA-seq dataset
- Efficient deep learning framework enables integration of millions of cells on a personal computer

## Introduction

The human body is composed of an estimated 30 trillion cells ^1^. The Human Cell Atlas (HCA) project ^2^ aims to create a comprehensive reference map of all cells of the human body. To accomplish this goal, the HCA leverages recent advancements in single cell technologies which enable the profiling of thousands to up to millions of cells in routine experiments ^3,4^. The HCA is a community-driven initiative that motivates research groups across the world to contribute. However, due to the minute amount of starting material, scRNA-seq data is prone to batch effects ^5^, which obstruct analysis of data generated by different laboratories, platforms, or experiments. Therefore, one of the most critical challenges for the success of the HCA and other cell atlasing projects is the integration of many large-scale scRNA-seq data generated by various groups in diverse experiments. As we anticipate an increase in the number of experiments and volume of cells profiled per experiment, the need for integration of multi-million transcriptome datasets will become increasingly common.

A number of methods have been developed to integrate scRNA-seq data across experiments, such as BBKNN, Scanorama, and Harmony ^5–9^. However, most of these methods are time-consuming and require large computational resources. Thus, these methods are essentially unavailable to researchers without access to high performance computing systems. To solve this issue, we introduce a deep learning approach entitled INSCT (“Insight”). Our algorithm generates integrated cellular embeddings, where transcriptionally similar cells from various batches map near each other overcoming differences between batches.

Deep learning approaches have demonstrated high performance and scalability in complex data ^10^. This is particularly true for image analysis ^11^ but the number of deep learning applications in molecular biology has been increasing steadily ^12,13^. Thanks to the large number of data points generated in a given experiment, scRNA-seq data has become attractive to the application of deep learning techniques ^4^. We ^14^ and others ^15,16^ have previously developed deep learning approaches tailored towards the analysis of scRNA-seq data.

INSCT is based on triplet neural networks (TNN) which have been primarily used in image recognition ^17^ and recently for unsupervised dimension reduction of scRNA-seq data ^18^. Here, we significantly extend traditional TNNs by formulating the novel concept of batch-awareness. By sampling triplets in a batch-aware manner, INSCT efficiently integrates up to millions of single transcriptomes.

To demonstrate validity and usage of INSCT, we apply our method to a number of simulated and real scRNA-seq datasets. Additionally, we benchmarked our method with respect to performance and scalability by comparison to existing state-of-the-art scRNA-seq integration methods. To enable broad usability, INSCT integrates seamlessly downstream of the popular scRNA-seq analysis framework Scanpy ^19^ and is freely available via our Github link.

## Results

TNNs learn data representations by distance comparisons ^20^. More precisely, triplets of data points are defined consisting of the Anchor, Positive and Negative inputs (Fig. 1). The loss function then minimizes the distance between the Anchor and Positive pair and maximizes the distance between the Anchor and Negative pair. Anchor and Positive are transcriptionally similar cells and the Negative is a cell transcriptionally dissimilar to the Anchor. We use K nearest neighbors (KNNs) and mutual nearest neighbors (MNNs) to define transcriptional similarity. To overcome batch effects, we introduce the concept of batch-aware triplet sampling. Triplets are sampled such that most Anchor-Positive pairs come from two different batches, while the Anchor-Negative pairs come from the same batch. The network learns a transformation which brings the Anchor-Positive pair closer to each other while pushing the Anchor-Negative pair further apart. Thus, the network learns a data representation that overcomes batch effects resulting in an integrated embedding.

**Fig. 1.**
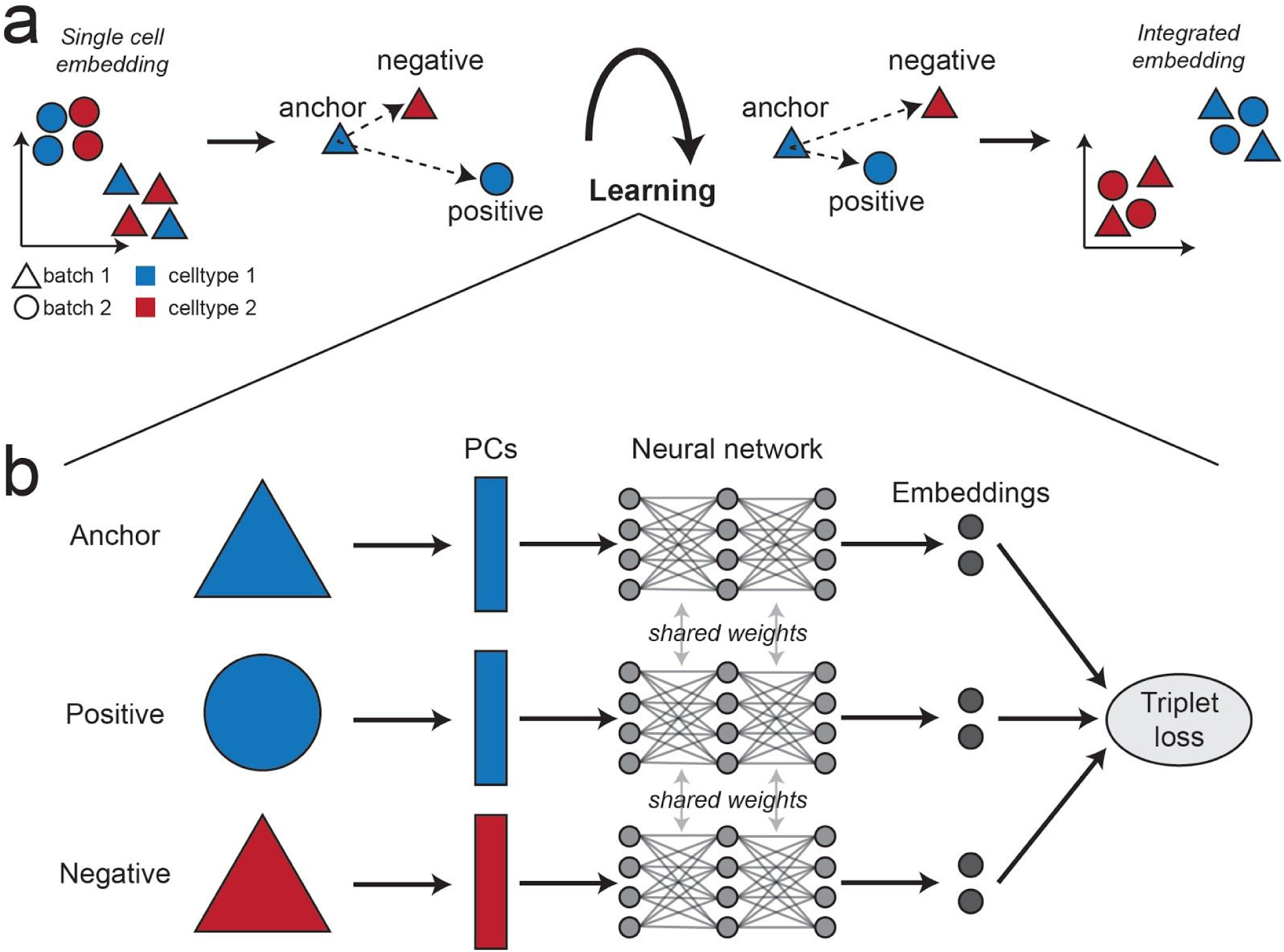
Overview of INSCT. **a**, INSCT learns a data representation, which integrates cells across batches. The goal of the network is to minimize the distance between Anchor and Positive while maximizing the distance between Anchor and Negative. Anchor and Positive pairs consist of transcriptionally similar cells from different batches. The Negative is a transcriptomically dissimilar cell sampled from the same batch as the Anchor. **b**, Principal components of three data points corresponding to Anchor, Positive and Negative are fed into three identical neural networks, which share weights. The triplet loss function is used to train the network weights and the two-dimensional embedding layer activations represent the integrated embedding.

### INSCT overcomes batch effects in scRNA-seq

As a proof of concept and to evaluate our method in a controlled setting, we applied INSCT to simulated data. Using the splatter ^21^ framework, we simulated two scRNA-seq specific scenarios. In scenario 1, four cell groups were generated from three batches. Application of standard UMAP or ivis dimension reduction resulted in 12 distinct clusters, indicating that the batch effect has not been removed (Fig. 2a). In contrast, the cellular embedding calculated by INSCT given the batch information as input, resulted in four distinct clusters corresponding to the four simulated cell groups, demonstrating that batch effects were successfully removed.

**Fig. 2.**
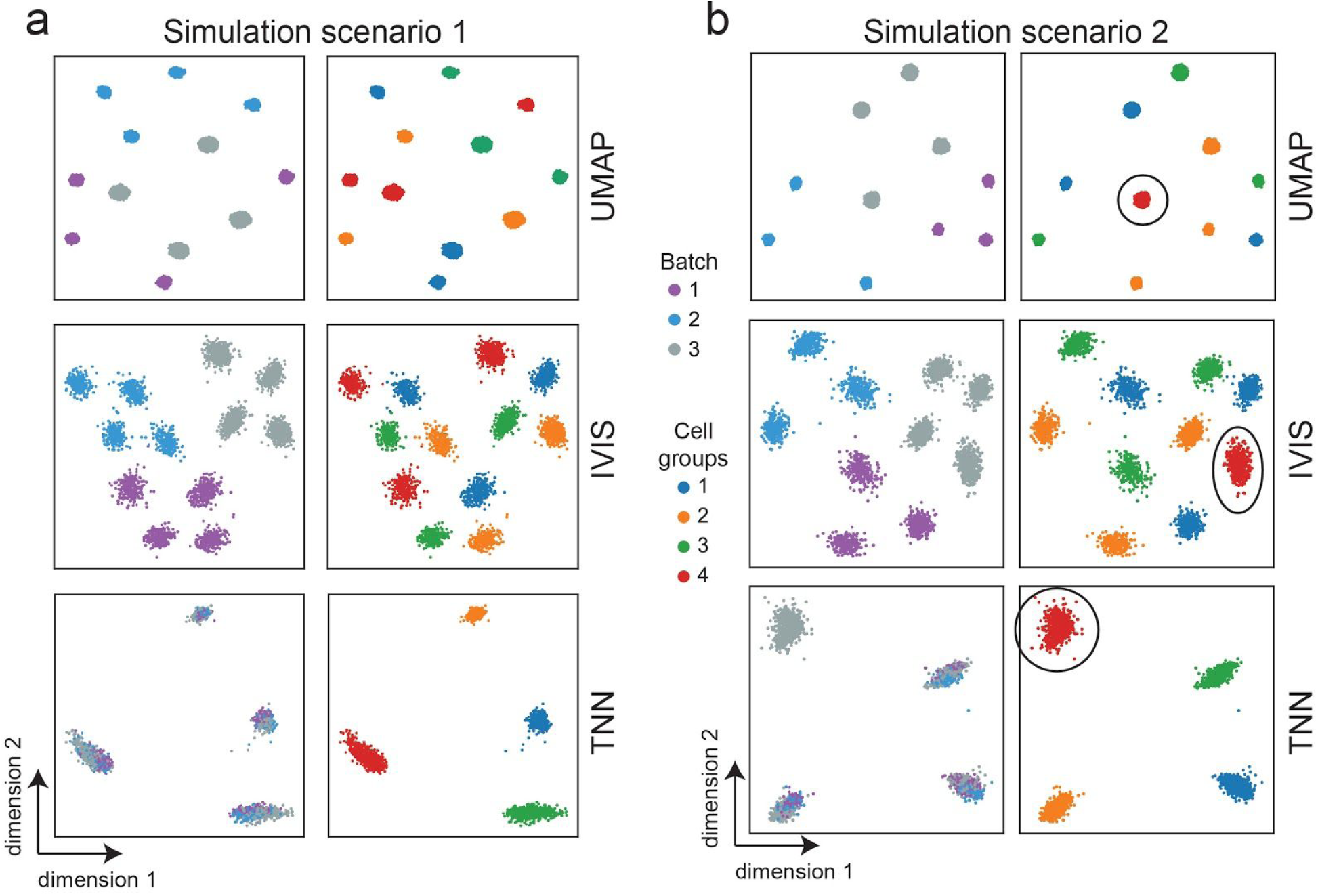
INSCT robustly overcomes batch effects in simulated scRNA-seq data. **a**, Reduced dimensions derived from UMAP (top), IVIS (middle) and INSCT (bottom) display cells from simulation scenario 1 consisting of three different batches and four cell groups. **b**, Reduced dimensions derived from UMAP (top), IVIS (middle) and INSCT (bottom) display cells from simulation scenario 2 consisting of three different batches and four cell groups of which one group is exclusively present in batch 3 (highlighted in black circles). For all panels, cells are colored by batch (left) and cell group (right).

Furthermore, to demonstrate that our algorithm does not “over-integrate” data by incorrectly mapping cells on top of each other across batches, we simulated scenario 2. Here, we removed one cell group (cell group four) from two batches such that cell group four was exclusively present in batch three. This group represents cells that are unique to a single batch and do not appear in any of the other batches. A situation that is frequently encountered in scRNA-seq analysis. Integration with INSCT maintained this group as a distinct cluster of cells (Fig. 2b), demonstrating that INSCT accurately integrates scRNA-seq data across batches and robustly preserves cells exclusively present in a single batch.

Since not all cells are guaranteed to have MNNs, it is important to contain sufficient KNN pairs in the training. Otherwise, the network will never include any cells without MNNs and, therefore, it will not learn a good transformation for these cells. At the same time, if the training does not include sufficient MNNs the batch effect will not be overcome completely. Therefore, we implemented the *k-to-m-ratio* parameter, which determines the number of Anchor-Positive pairs that are sampled based on KNN relative to MNN pairs. In Supplemental Figure 1 we demonstrate the robustness of INSCT with respect to a large range of the *k-to-m-ratio* parameter and how this parameter affects the integrated embedding.

### INSCT overcomes batch effects in real data

Next, we applied INSCT to real scRNA-seq data. To evaluate and benchmark INSCT, we analyzed a scRNA-seq data collection from the Tabula Muris project ^22^. The authors used two distinct technical approaches to profile nearly 100,000 individual cells from 20 mouse organs and tissues. The data consists of FACS-based scRNA-seq (FACS) experiments with 54,837 cells and droplet-based scRNA-seq (Droplet) experiments with 42,192 cells, respectively. Despite the fact that both technologies profiled identical tissues and cell types from the same animals, standard dimension reduction revealed minimal overlap between these two technologies (Fig. 3a). Moreover, when coloring the cells by the harmonized cell type ontology labels provided in the original publication, most cell types formed two distinct clusters separated by technology (Fig. 3b). After application of INSCT, cells from both technologies annotated to the same cell ontology mapped into corresponding areas of the integrated embedding for both FACS (Fig. 3c) and Droplet (Fig. 3d). The integrated embedding showed strong overlap between both datasets (Fig. 3e).

**Fig. 3.**
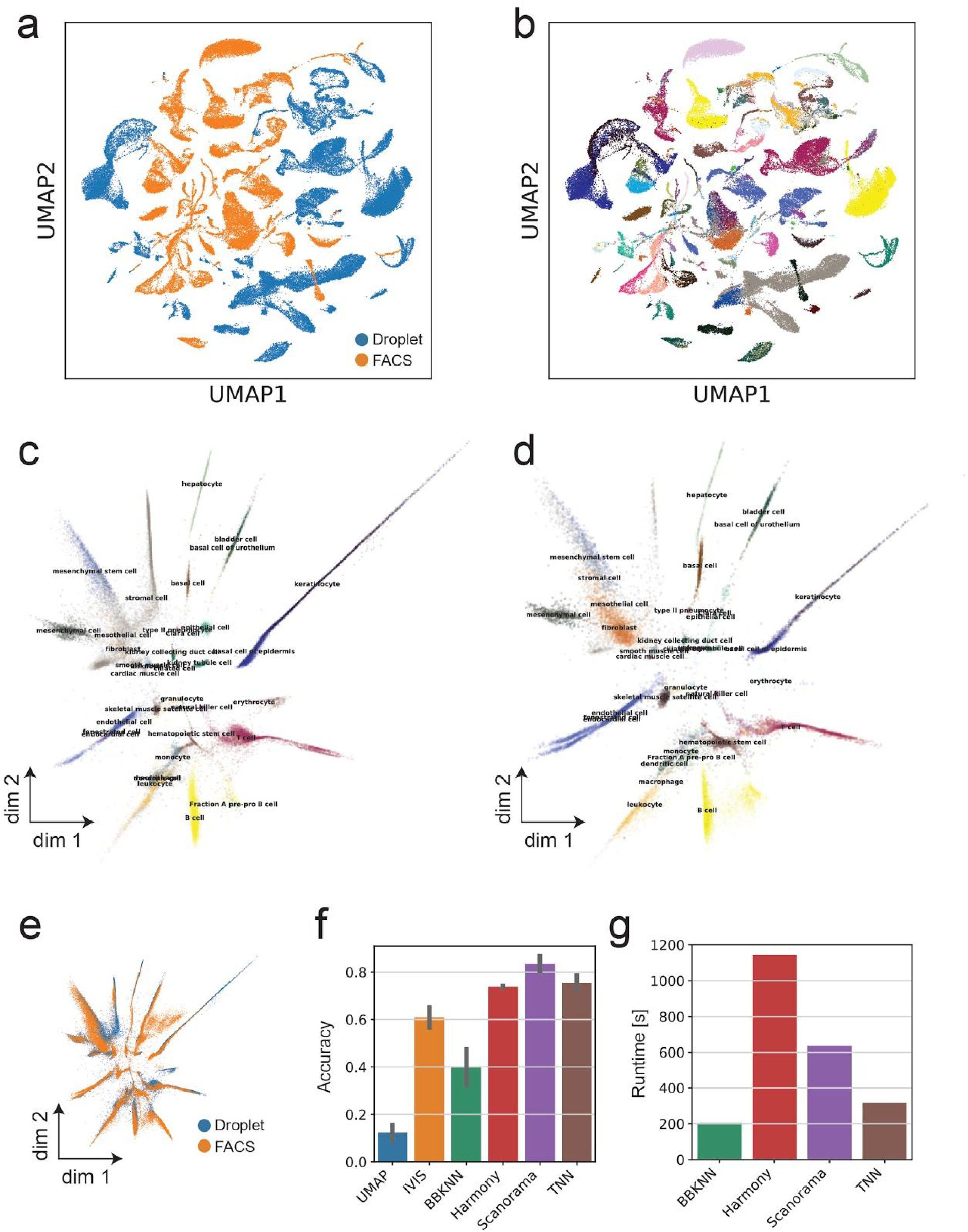
INSCT accurately integrates cells across different platforms. Standard UMAP embedding is depicted. Cells are colored by experiment technology (**a**) and cell ontology labels provided by the original publication (**b**). Integrated embedding of cells in both Droplet (**c**) and FACS (**d**) based scRNA-seq data is displayed. Cells are colored by cell ontology class as provided in the original publication. **e**, Scatterplot shows the integrated embedding of all cells. Cells are colored by platform. Barplots depict integration accuracy (**f**) and runtime (**g**) of various methods.

To assess the integration performance, we compared INSCT to three recent integration methods Harmony ^23^, Scanorama ^7^ and BBKNN ^8^. We selected these three algorithms because they showed high scalability in previous comparisons ^24^ and are written in Python. Next, we applied each of these three methods to integrate the Tabula Muris data collection (Fig. S2).

To quantitatively evaluate the integration, we used the following approach. First, we trained a Nearest Neighbor classifier on the integrated embedding coordinates of one of the two batches to predict the cell type annotations. Next, we predicted the cell type annotations for the left out batch and compared the predicted to the original cell type annotations. The rationale behind this approach is that cells of the same type should fall close to each other in the integrated embedding space regardless of batch. If corresponding cell types were mapped near each other in the integrated embedding, then the classifier would achieve high accuracy. We used the accuracy of these classifiers as a proxy for the integration performance. INSCT demonstrated high performance compared to competing methods (Fig. 3f). Since this dataset consisted of nearly 100,000 cells, we also compared runtimes of the different methods. Regarding runtime, INSCT ranked among the fastest methods and finished integration in under five minutes (Fig. 3g). To further demonstrate the robustness of our method, we applied INSCT to a second independent collection of scRNA-seq data from human pancreas (Fig. S3).

### INSCT robustly integrates cells across species

Next, we set out to test whether INSCT can be used to integrate cell types across species. We downloaded scRNA-seq data from human ^25^ and mouse ^26^ lung cells. The hypothesis was that cells of the same type should map into corresponding regions of the integrated embedding space. Therefore, we applied INSCT to generate an integrated embedding after combining the human and mouse data (Fig. 4a). Overlaying the corresponding cell type annotations demonstrated that both human and mouse cell types mapped into the same regions of the embedding (Fig. 4bc). For example, human and mouse alveolar type 1 and 2 cells mapped into the same region of the integrated embedding. Additionally, cell type marker genes showed homologous expression patterns for both human and mouse cells (Fig. 4d). To assess the correspondence between human and mouse cells, a random forest model was trained on the mouse cell embedding coordinates and subsequently used to predict the cell types of the human cells. The majority of predicted and observed cell types showed strong agreement between human and mouse harmonized cell types (Fig. 4e). This analysis demonstrated that homologous human and mouse cell types map into the same region in the integrated embedding. Therefore, INSCT can robustly integrate cells across species.

**Fig. 4.**
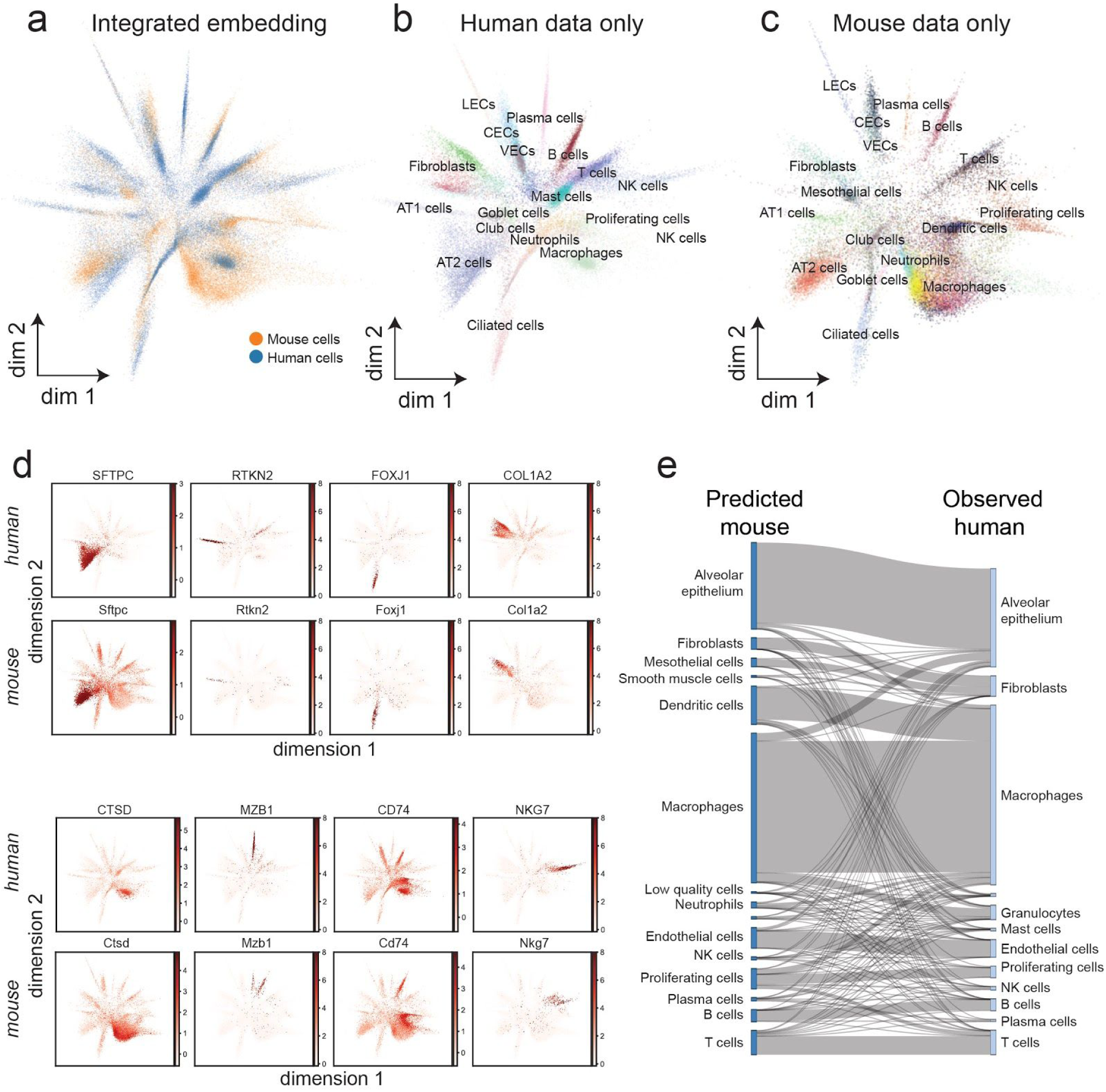
INSCT enables cross-species integration. Integrated embedding space represents cells from both species (**a**), human (**b**) and mouse (**c**) cells only. The colors correspond to cluster annotations taken from the original publications. CECs = capillary endothelial cells, LEC = lymphatic endothelial cells, VECs = vascular endothelial cells. **d**, Expression of cell type marker homologs in both human and mouse cells across the integrated embedding. Red and white colors represent high and low relative expression values, respectively. **e**, Sankey diagram illustrates correspondence between predicted mouse cell type labels and observed human cell types.

### Semi-supervised INSCT robustly classifies cell types

So far, we have applied INSCT in an unsupervised manner. However, when cell annotations are available, this information can be incorporated into the model. Therefore, we extended the functionality of INSCT by sampling Anchor-Positive pairs based on available cell type labels in addition to MNNs or KNNs. To demonstrate and explore the effects of this additional functionality, we analyzed scRNA-seq data collection from four independent scRNA-seq studies of human pancreas cells. Each study used a different technology to profile single cells. As expected, training INSCT in a supervised mode increased separation between cell types (Silhouette coefficient=0.7572, Fig. 5a) compared to completely unsupervised mode (Silhouette coefficient=0.2061, Fig. S3).

**Fig 5.**
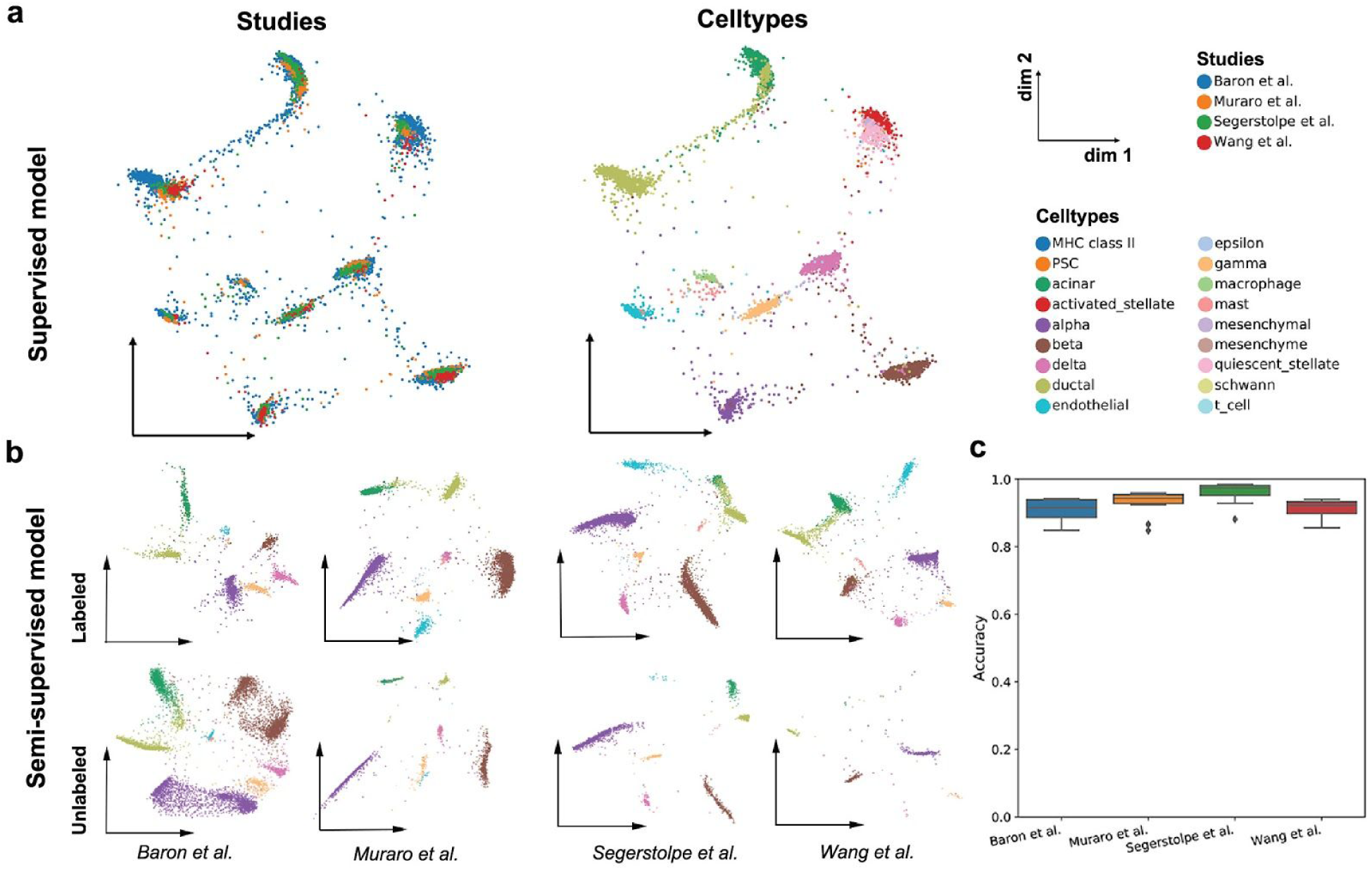
Semi-supervised INSCT robustly classifies cell types. **a**, Integrated embedding plots generated from supervised INSCT display separation of cell types. Cells are colored by study (left) and cell type (right). **b**, Integrated embedding plots are displayed for four independent semi-supervised INSCT runs. In each run the cell type labels from one study were masked. From left to right the masked studies were Baron et al., Muraro et al., Segerstople et al. and Wang et al. respectively. Integrated embedding for the three labeled and each unlabeled study are depicted on top and bottom, respectively. Colors correspond to common cell types. **c**, Boxplots display prediction accuracies (y-axis) from ten bootstrapped runs for each masked study.

Next, we set out to investigate the use of INSCT in a semi-supervised mode. Semi-supervision can be applied when part of the data has missing labels. In this situation the triplet sampling will be based on 1) the cell type labels between the labeled studies and 2) MNNs calculated from the studies without labels. Thus, the network integrates both information from the cell type labels as well as transcriptional similarities. Semi-supervision enables users to integrate an unlabeled target scRNA-seq data set into a labeled reference collection. We iteratively trained INSCT based on a leave-one-out approach. In each iteration, the cell type labels from three studies were used and the labels from the fourth study were masked during training. The integrated embeddings showed clear separation between cell types and good overlap across studies (Fig. 5b). Moreover, a classifier trained on the integrated embedding coordinates achieved high accuracies across all four iterations (Fig. 5c). Since manual cell type annotations can be error-prone or different in labeling, we explored the impact of incorrect labeling using our simulated data. Indeed, since INSCT uses both information from the cell type labels as well as transcriptional similarities, INSCT is robust to the incorrect labeling (Fig. S4). Together these findings illustrate the ability of INSCT to correctly integrate unlabeled and labeled scRNA-seq data.

### INSCT scales to multi-million brain transcriptomes

To showcase the efficiency of our algorithm we attempted to integrate over 2.6 million mouse brain transcriptomes from four independent resources. To the best of our knowledge, this represents the largest scRNA-seq integration that has been reported to date. Data was collected from the following sources: 1) 10x Genomics (n = 1,292,016), 2) Dropviz (n = 691,600), 3) SPLIT-seq (n = 156,049), and 4) the MouseBrainAtlas (n = 504,020). All 2.6 million transcriptomes were merged into one object and standard preprocessing was performed consisting of identification of highly variable genes, normalization and initial dimension reduction by principal component analysis. The resulting principal components were used as input into INSCT, which was run on a freely available Google Colab instance in a standard web browser. The model converged in less than 1.5 hours and the resulting integrated embedding showed good overlap between all four datasets (Fig. 6a).

**Fig. 6.**
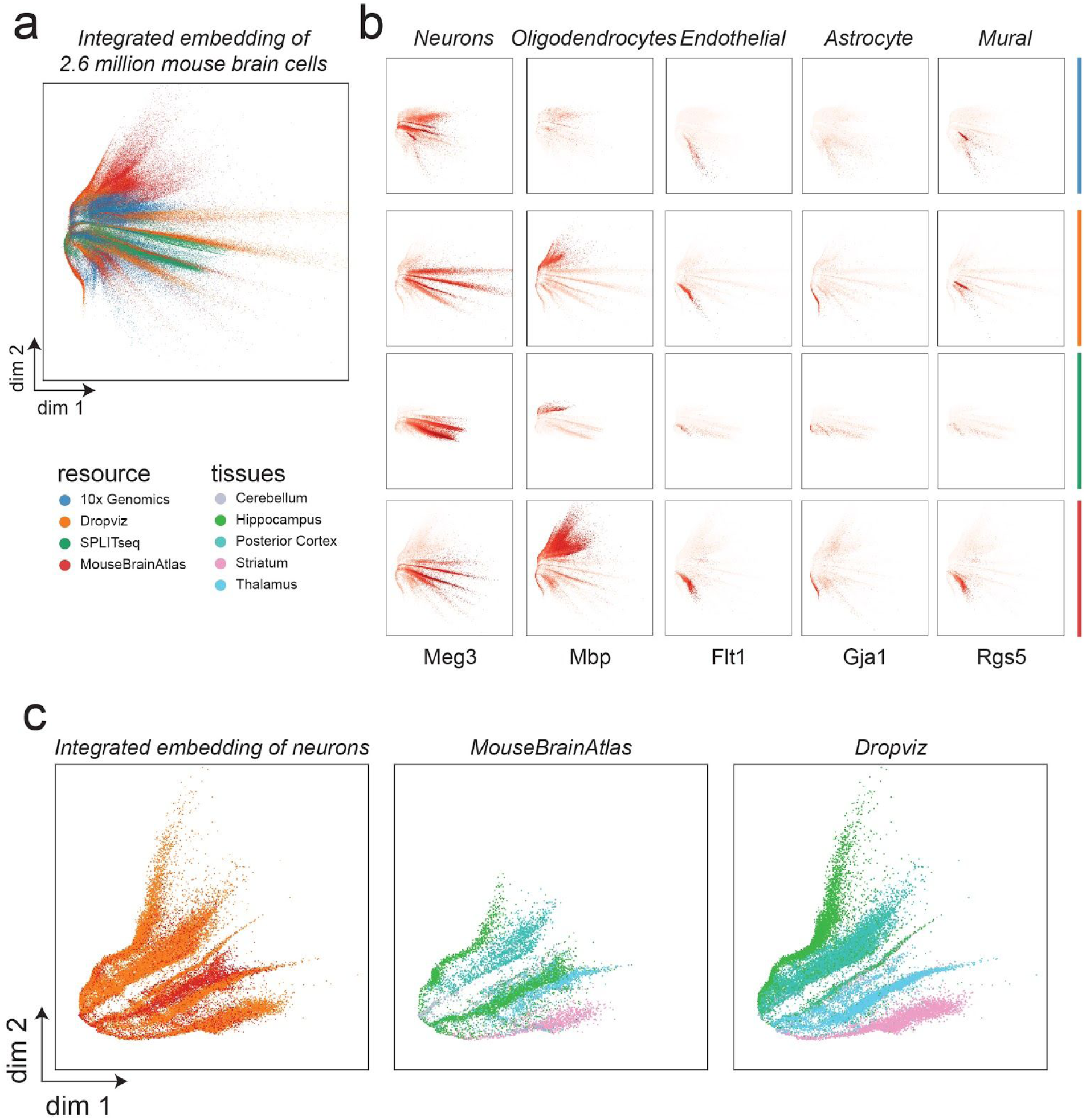
INSCT integrates multi-million brain cells. **a**, Integrated embedding displays 2.6 million brain transcriptomes. Cells/nuclei are colored by the originating study. **b**, Expression patterns of maker genes across all for studies in the integrated embedding are shown. Cells are split by studies (rows) and cell type markers (columns). **c**, Integrated embedding of neurons from both the Dropviz and MouseBrainAtlas are depicted for cells from Cerebellum, Hippocampus, Posterior Cortex, Striatum and Thalamus (left). Cells are colored by study. The same embedding space is shown for MouseBrainAtlas (middle) and Dropviz (right) cells only. Colors represent brain regions.

Specific cell types, characterized by expression of marker genes, were mapped into corresponding regions of the integrated embedding across all four resources (Fig. 6b). Expression of marker genes *Gja1, Mbp, Flt1* and *Rgs5* for cell types astrocytes, oligodendrocytes, endothelial and mural cells, respectively, overlapped across all four sources. The lack of clear expression patterns of all of these genes in the 10x Genomics and SPLIT-seq data sets indicated that these cell types appeared at a relatively low frequency in both of these two datasets. A large number of cells were marked by the expression of *Meg3*, a well known neuron marker. The neurons from all four datasets were mapped into the same embedding space, while still showing neuron-specific heterogeneity.

To validate the accuracy of the integrated embedding space, we focused on the neurons, which are known to display high levels of transcriptional heterogeneity ^27^. Therefore, we restricted the data to neurons and ran INSCT on this subset. Both the Dropviz and MouseBrainAtlas data contained annotations indicating which brain region had been sampled. We observed good overlap between neurons derived from corresponding brain regions in these two datasets. For example, neurons from the Thalamus, Straitum and Cerebellum from both datasets were mapped into the corresponding regions of the integrated embedding (Fig. 6c). Collectively, these results demonstrate that INSCT accurately and efficiently integrates multi-million transcriptome datasets.

### INSCT efficiently integrates millions of cells

To quantitatively assess scalability and efficiency of the integration methods, we used our collection of 2.6 million mouse brain transcriptomes. The merged dataset was down-sampled to varying proportions and subsequently integrated using BBKNN, Harmony, Scanorama and INSCT. For each integration, the runtime was measured (Fig. 7). Our analysis indicated that INSCT provides substantial speed advantage for data sets of more than 100,000 cells. Runtime benchmarking was conducted on a Google Colab instance restricted to a maximum of 25 gigabytes of memory and two cores. BBKNN and Scanorama ran out of memory at 400,000 and 1,600,000 cells, respectively and therefore did not complete. While both BBKNN and Scanorama are among the most scalable existing integration tools ^24^, these results indicate that both methods require large computational resources for integrations of this order of magnitude.

**Fig. 7.**
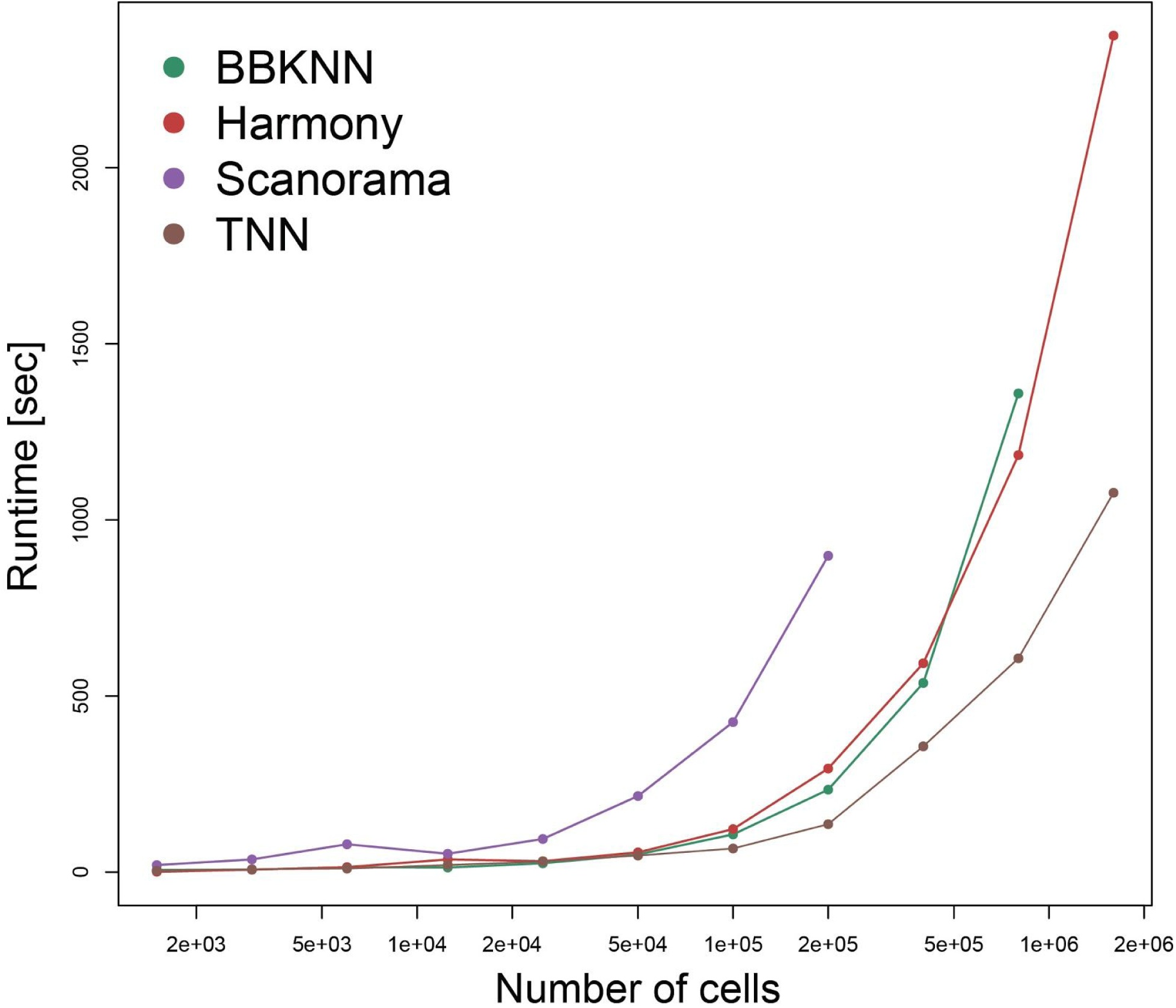
INSCT efficiently integrates millions of cells. The graph shows the number of cells and their runtime in seconds on the X and Y axes, respectively. Colors represent the different integration methods.

## Discussion

Here, we introduced a novel deep learning method for the integration of scRNA-seq data. We note that INSCT achieves competitive accuracies compared to current state-of-the-art scRNA-seq integration methods. However, with respect to scalability and efficiency INSCT significantly outperforms all other methods. This represents an important advancement, as it empowers researchers without access to high performance computing systems to perform integration of millions of transcriptomes. Additionally, in contrast to existing integration methods, INSCT can robustly project unseen data into pre-generated embeddings. Therefore, a down-sampling strategy can be used to further decrease runtime by training the network on a subset of cells and subsequently projecting the remaining cells into the embedding space.

In our analysis we showed that INSCT accurately integrates cells across species. This approach can be applied when homologous cell types are not known *a priori* or one of the experiments does not contain cell annotations. Thus, we believe that INSCT is a valuable tool for cross-species single transcriptome studies.

As the number and density of cell atlases increases, we expect more and more researchers to project their primary scRNA-seq data into the context of these annotated reference atlases. INSCT in semi-supervised mode enables users to robustly integrate and classify their unlabeled primary data into well-annotated reference data collections.

While we applied INSCT exclusively to scRNA-seq data, in principle, our network can be applied to other data types. In the future we plan to extend the application of INSCT to other data types, including imaging data, where batch effects obstruct integrated analysis.

Finally, all analyses described in this manuscript are available via interactive Google Colab notebooks. These notebooks allow anybody using a common web browser, such as Google Chrome, to reproduce analyses and figures interactively. Users can explore INSCT and our analyses by changing any parameter or substituting the underlying data. INSCT source code and interactive usage tutorial is freely available at https://github.com/lkmklsmn/insct.

## Methods

INSCT makes good use of the source code of ivis ^18^ (https://github.com/beringresearch/ivis) and was written in Python. Moreover, INSCT is designed to seamlessly integrate downstream of the popular scRNA-seq analysis framework Scanpy ^19^.

### Batch-aware triplet sampling

Each triplet is formed by defining an Anchor, a Positive and a Negative cell. The Positive is transcriptionally similar to the Anchor, while the Negative is transcriptionally dissimilar to the Anchor. For batch-aware triplet sampling, we make use of K nearest neighbors (KNNs) and mutual nearest neighbors (MNNs) to define transcriptionally similar cells. MNNs were previously introduced for batch correction and build the foundation of many scRNA-seq integration tools ^5^.

Positive cells are defined in a two-step approach. First, MNNs are calculated between all possible batch pairs. For any given Anchor, all MNNs are defined as Positives. This procedure ensures that Anchor-Positive pairs are sampled from two different batches and thereby forces the network to overcome the batch effect. Depending on the overlap of cell types between batches, only a fraction of cells typically fall into the MNN category. To inform the network of the remaining data manifold, we calculate KNNs across all cells for a subset of residual cells. This ensures the network to learn a data representation for cell types that are exclusively present in one batch.

Negatives are cells randomly sampled from the same batch as the Anchor. The K parameter defines the number of nearest neighbors and is usually much smaller than the total number of cells. Therefore, the likelihood of erroneously drawing a cell from within the K nearest neighbors is very small and a randomly sampled cell is highly likely to be transcriptionally dissimilar to the Anchor. Thus, random sampling provides a fast and practical way to define Negatives.

For supervised and semi-supervised training, triplets are sampled based on the cell type annotations in addition to MNNs and KNNs. The label_ratio parameter controls the number of triplets that are sampled from the cell type annotations and is defined as the proportion of triplets sampled from all labeled cells. Inclusion of triplets sampled from MNNs and KNNs ensures that the training is not completely dependent on the cell annotations. As a consequence, the network is robust to errors in the cell annotations as demonstrated in Supplemental Figure 4.

INSCT implements two different KNN search algorithms. Users can perform exact KNN search as implemented in the Python *sklearn* package. Alternatively, INSCT provides an efficient KNN approximation as implemented in the Python *hwnslib* package. By default, INSCT uses the KNN approximation which is highly recommended for large datasets.

### Network architecture and training

The network architecture was derived from ivis with following adaptations. The base network consists of three layers with dimensions defined as 75% of the dimension of principal components that were used as inputs. Each of these dense layers is interleaved by Alpha Dropout layers with a dropout rate of 0.25. The input principal components of three data points corresponding to Anchor, Positive and Negative are fed into three identical base networks, which share their weights and are connected to a two-dimensional embedding layer. The activations of the embedding layer represent the integrated embedding.

After having computed the Positives for each Anchor, triplets are generated dynamically during training. At the beginning of each training epoch, a single triplet combination is sampled for each Anchor. Given the number of cells profiled in any scRNA-seq experiment, the likelihood of sampling identical triplets is extremely small. Therefore, the network is ensured to be presented with different sets of triplets in each epoch.

Furthermore, the triplet loss is optimized by minimizing the distance between Anchor-Positive pairs and maximizing the distance between Anchor/Positive-Negative pairs. Here, the triplet loss was defined as follows:

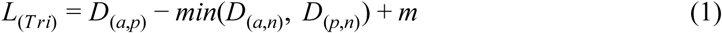

where *a, p* and *n* correspond to Anchor, Positive and Negative respectively. *D*_(·,·)_ is defined as euclidean distance, and m is a margin between similar and dissimilar pairs. Training is automatically ended once the loss does not improve over a specified number of epochs.

### Data sources

#### Simulation data

Simulations were performed using the splatter framework ^21^. For simulation scenario 1, 2000 genes and three batches with 1000, 1000 and 2000 cells were simulated using the following parameters: batch.facLoc = 0.3, batch.facScale = 0.3. Additionally, four cell groups with equal probabilities were simulated in each batch. For scenario 2, all cells from group four were removed from batches one and two so that cell group four was exclusively present in batch three.

#### Pancreas data

Pancreas data collection consisted of four independent studies using four different technologies, including CEL-seq (GSE84133) ^28^, CEL-seq2 (GSE85241) ^29^, SMART-seq (E-MTAB-5061) ^30^, and Fluidigm C1 (GSE83139) ^31^. Each dataset was subsequently merged resulting in a Scanpy AnnData object consisting totaling 14,844 cells.

#### Tabula Muris data

The Tabula Muris mouse atlas expression data was downloaded from (https://tabula-muris.ds.czbiohub.org/), which contained nearly 100,000 cells from 20 organs and tissues. This dataset consisted of two distinct technical approaches, including FACS-based scRNA-seq experiments with 54,837 cells and droplet-based scRNA-seq experiments with 42,192 cells, respectively.

#### Lung data

Mouse lung scRNA-seq data consisted of 29,297 cells and was obtained from https://github.com/theislab/2019_Strunz. The human lung scRNA-seq data consisted of 41,889 cells and was obtained from https://github.com/theislab/2020_Mayr. Cell type annotations were taken from corresponding Github repositories.

#### Mouse brain data

Mouse brain single cell/nuclei RNA sequencing data was downloaded from four different resources. The 10x Genomics dataset consists of 1,306,127 mouse brain cells and was downloaded from https://support.10xgenomics.com/single-cell-gene-expression/datasets/1.3.0/1M_neurons. The DropViz dataset containing 691,600 cells from nine different mouse brain regions was downloaded from http://dropviz.org. The MouseBrain.org data was downloaded as separate tissue-level files from http://mousebrain.org/downloads.html consisting of a total of 507,287 cells. The SPLIT-seq data^32^ consisting of 156,049 single-nucleus transcriptomes from postnatal day 2 and 11 mouse brains and spinal cords was downloaded from the Gene Expression Omnibus (Accession: GSE110823).

### Data processing

All data preprocessing was performed using the Python package Scanpy. Raw count expression matrices were imported and merged into a single Scanpy AnnData object using the Scanpy *concatenate* function. The merged object was further filtered by requiring that at least 200 genes were detected per cell and each cell contained less than 10% mitochondrial reads. Data was subsequently normalized using the *normalize_per_total* function. Genes observed to be highly variable within a data set were calculated using the *filter_genes_dispersion* function. Next, the object was scaled and subjected to principal component analysis. The resulting principal components were used as the inputs into INSCT.

### Integration accuracy

To evaluate the performance of the integration procedure, we applied the following approach. To assess the integration performance, we trained a Nearest Neighbor classifier as implemented in the *sklearn* module using the coordinates of the integrated embedding as the input to predict the cell annotations. The implementation of the classifier was taken from the sklearn Python package. The classifier was trained on one or more batches and subsequently used to predict the cell annotations of the left out batch. The accuracy derived from comparison between the predicted and observed cell annotations was used as an indicator of integration performance.

### External software

For evaluation purposes we compared INSCT to existing scRNA-seq dimension reduction and integration tools. The evaluation included UMAP (version 0.3.10), ivis (version 1.6.0), Harmonypy (version 0.0.4), Scanorama (version 1.5), and BBKNN (version 1.3.6).

### Computing environment

All integration analyses were performed in a freely available Google Colab instance, which is limited to two cores and a maximum of 25GB memory. The code to interactively reproduce the results described in this manuscript can be run by anybody with access to a web browser and without any access to high performing computing systems or computers.

## Authors’ contributions

LMS conceptualized the algorithm. LMS, YYW and ZZ designed the project. LMS and YYW developed the algorithm and analyzed the data. ZZ supervised the project. LMS, YYW, and ZZ wrote the manuscript. All authors read and approved the final manuscript.

## Acknowledgments

The authors would like to thank the members of the Bioinformatics and Systems Medicine Laboratory at the University of Texas Health Science Center at Houston as well as David Henke for stimulating discussion. This work was supported by the Cancer Prevention and Research Institute of Texas (CPRIT core grant RP180734). ZZ was also partially supported by National Institutes of Health grant (R01LM011177). The funders had no role in the study design, data collection, and analysis, decision to publish, or preparation of the manuscript.

## Supplemental Figures

**Supplemental Figure 1.**
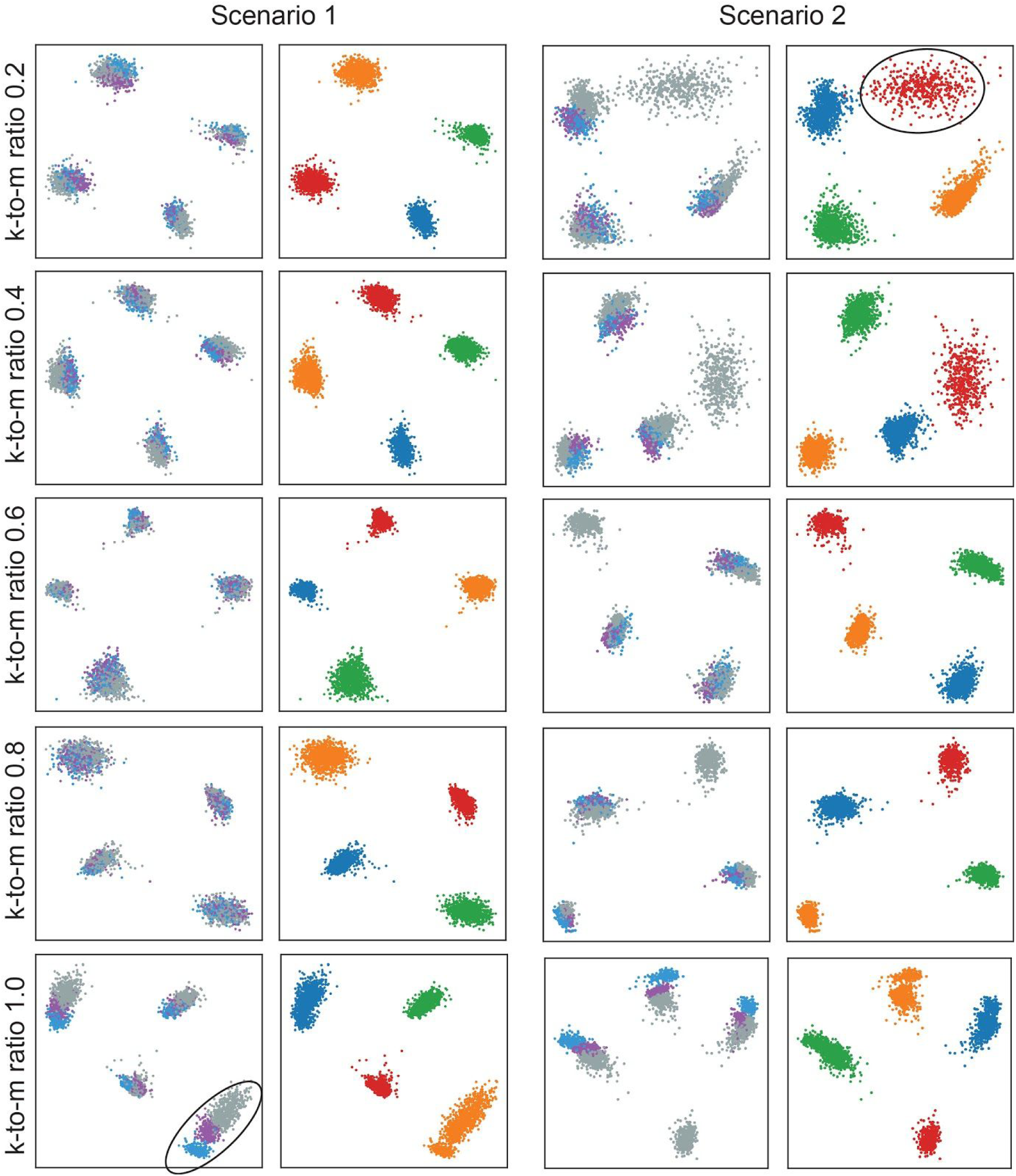
Embedding plots illustrate the robustness of the k-to-m ratio parameter in the simulation Scenarios 1 (left) and 2 (right). The k-to-m parameter determines the number of triplets sampled from KNNs relative to MNNs and increases from top to bottom. In scenario 1, the higher the k-to-m ratio parameter the stronger the within cluster batch effect (as highlighted in black ellipse). In scenario 2, the lower the k-to-m ratio parameter the less coherent is the cluster of cells exclusive to batch 3 (as highlighted in black ellipse). Since these cells do not have any MNNs the network can not learn a robust representation of these cells.

**Supplemental Figure 2.**
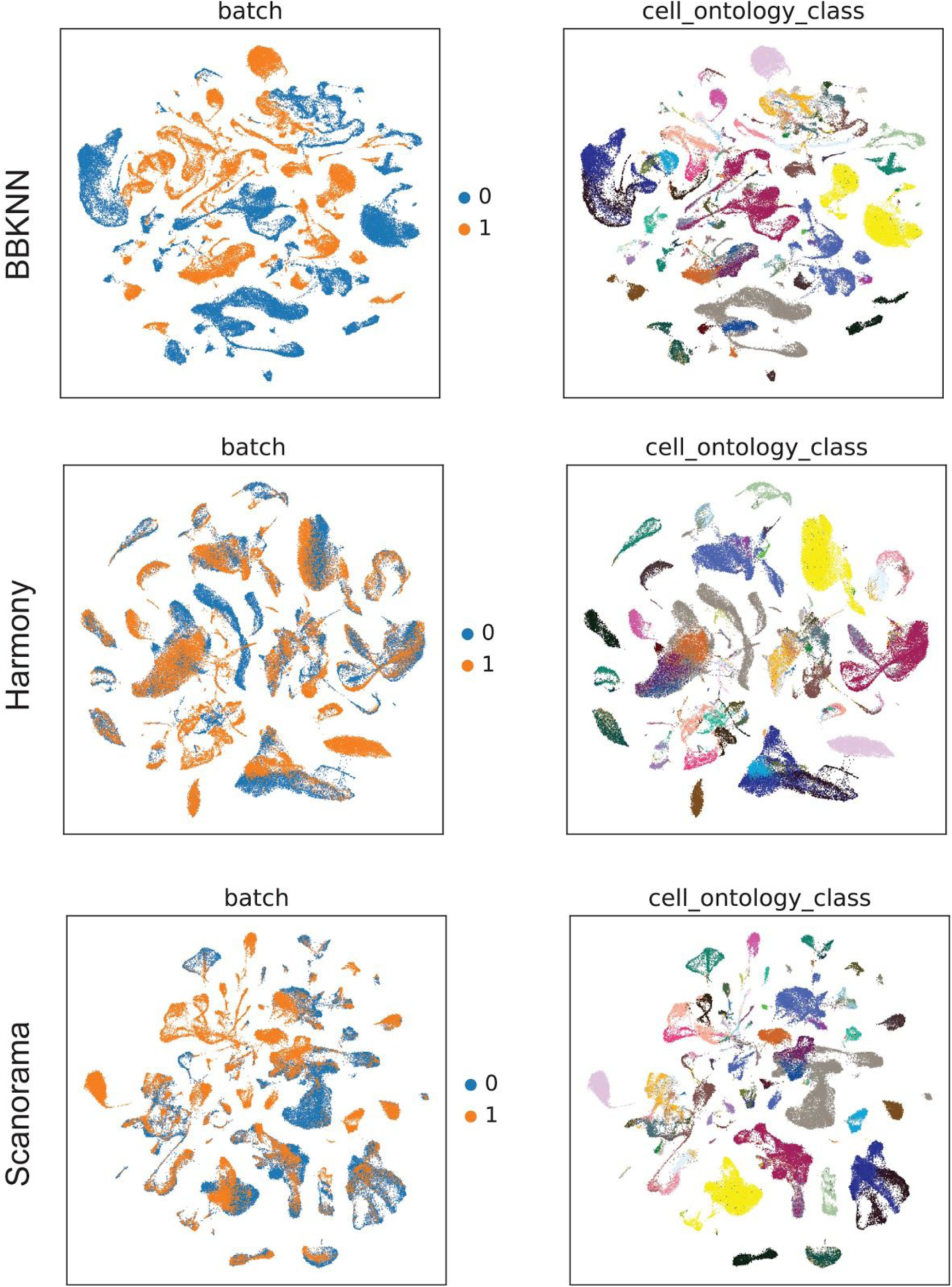
Integrated embedding plots for BBKNN (top), Harmony (middle) and Scanorama (bottom) applied to the Tabula Muris data. Cells are colored by batch (left) and cell type ontology (right).

**Supplemental Figure 3.**
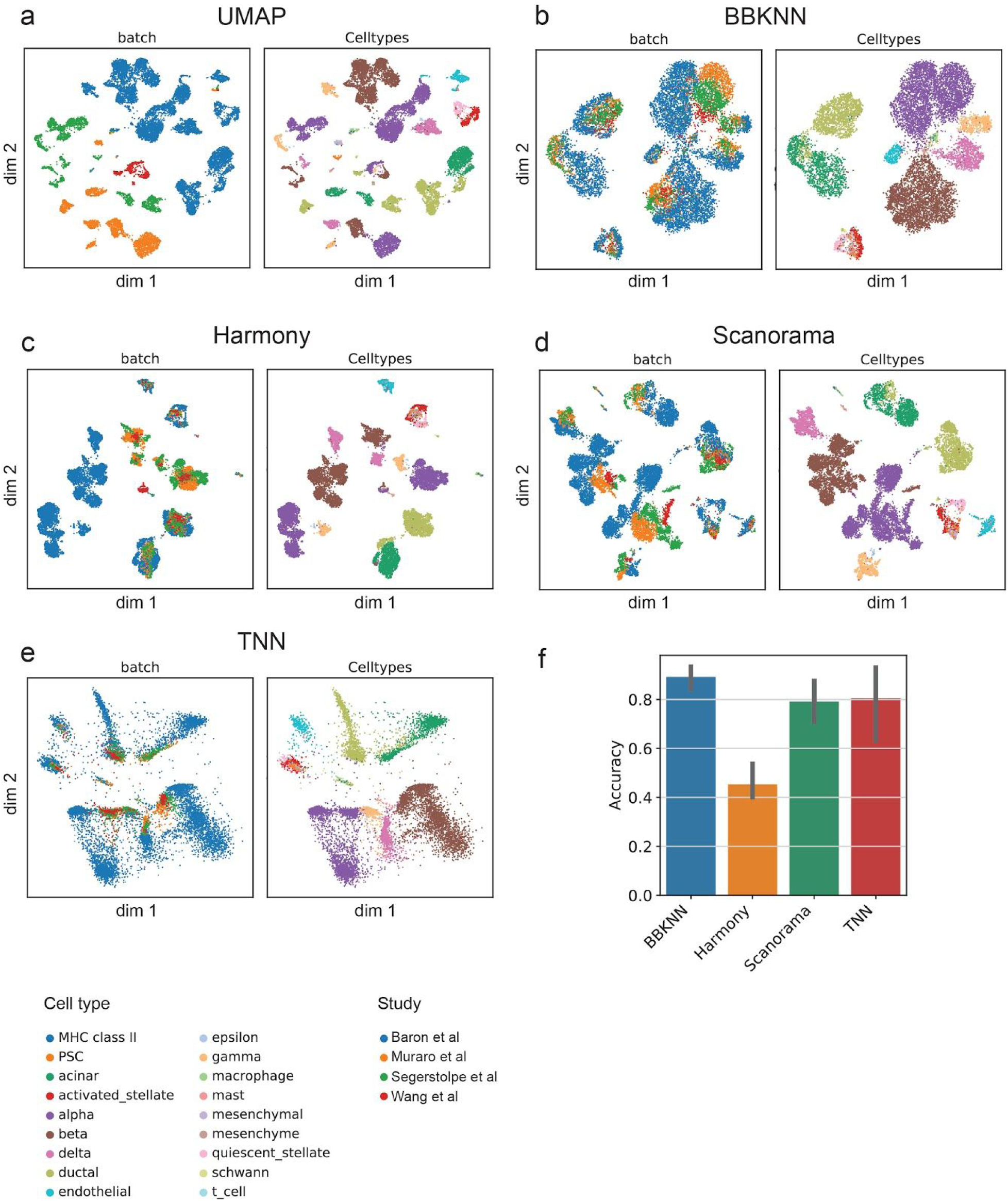
Embedding plots for UMAP (**a**), BBKNN (**b**), Harmony (**c**), Scanorama (**d**) and INSCT (**e**) applied to the Pancreas data collection. For each panel, cells are colored by batch (right) and cell type (left). (**f**) Random forest integration accuracies from the leave-one-out iterations across the four integration methods.

**Supplemental Figure 4.**
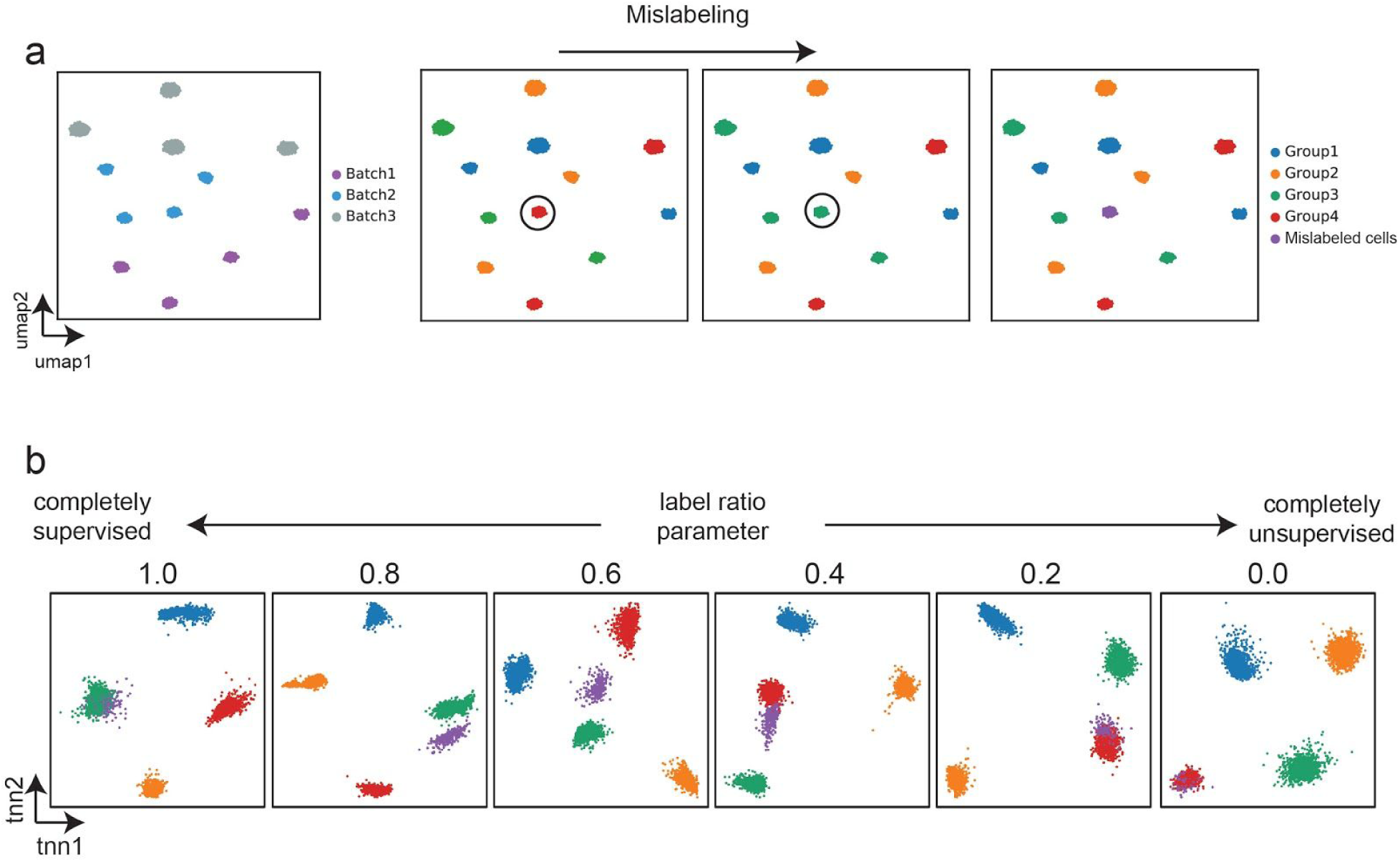
**a**, UMAP depicts cells from simulation scenario 1. Group of red cells from batch 2 is mislabeled as green cells. For visualization purposes only, the mislabeled cells are colored purple. **b**, As the label ratio parameter decreases from left to right the model training contains more triplets sampled from MNNs and KNNs compared to cell types, thereby going from completely supervised (label ratio 1) to completely unsupervised (label ratio 0). At high label ratio the mislabeled cells (purple) map onto the incorrect cluster (green). As the label ratio parameter decreases the mislabelled cells (purple) form a separate cluster. At label ratio 0, which corresponds to completely unsupervised training the mislabeled cells map to the correct cluster (red).

